# Choice-Induced Preference Change under a Sequential Sampling Model Framework

**DOI:** 10.1101/2025.10.23.684256

**Authors:** Douglas G. Lee, Giovanni Pezzulo

## Abstract

Sequential sampling models of choice, such as the drift-diffusion model (DDM), are frequently fit to empirical data to account for a variety of effects related to accuracy/consistency, response time (RT), and sometimes confidence. However, no model in this class has been shown to account for the phenomenon known as *choice-induced preference change*, wherein decision makers tend to rate options higher after they choose them and lower after they reject them (and often choose the option that they had initially rated lower). Studies have reported choice-induced preference change for many decades, and the principal findings are robust. The resulting *spreading of alternatives* (SoA) in terms of their subjective value ratings is incompatible with traditional sequential sampling models, which consider the rated values of the options to be stationary throughout choice deliberation. Here, we propose that extending the basic DDM to incorporate independent attributes into the drift rate can allow this class of model to account for SoA. Critically, the extended model assumes that choice deliberation does not always involve the same set of attributes as is involved in individual evaluations of the same options. We show that this model can generate SoA (while simultaneously accounting for consistency and RT), as well as the relationships between SoA and choice difficulty, attribute disparity, and RT previously reported in the literature.

## Introduction

### Established patterns of choice behavior

Value-based decision-making research often focuses on choices between two options, using subjective value ratings of the individual options to explain choice behavior. Many robust patterns have been reported with respect to both choice responses and response times (RT). A prime example is that difficult decisions (those between options of similar subjective value) yield choices that are generally less predictable (from ratings) and slower, relative to easy decisions (Milosavljevic et al., 2010; Ratcliff & McKoon, 2008). Decisions are also thought to be more difficult when the value estimates of the choice options are less certain, and such decisions also yield choices that are less predictable and slower (Gwinn & Krajbich, 2020; Lee & Coricelli, 2020; Lee & Daunizeau, 2020, 2021; Lee & Hare, 2023b). Apart from difficulty, decisions that involve more valuable options yield choices that are usually faster and often more predictable (Hunt et al., 2012; Polanía et al., 2014; Sepulveda et al., 2020; Shevlin et al., 2022; Shevlin &Krajbich, 2021; S. M. Smith & Krajbich, 2019; Teodorescu et al., 2016). Looking beyond overall value ratings, choices between options with high attribute disparity (e.g., where each option dominates on a different attribute) are faster (Bhatia & Mullett, 2018; Lee & Hare, 2023b; Lee & Holyoak, 2021). Each of these independent variables also positively relates to choice confidence (Brus et al., 2021; Folke et al., 2016; Lee & Hare, 2023b)).

### Choice-induced preference change

Beyond the variables commonly measured in experimental choice paradigms (outlined above), the phenomenon known as *choice-induced preference change* was first reported long ago (Bem, 1967; Brehm, 1956; Festinger, 1957; Harmon-Jones & Mills, 2019) and more recently demonstrated in a wide array of value-based decision-making studies (Alós-Ferrer et al., 2012; Chammat et al., 2017; Colosio et al., 2017; Coppin et al., 2010, 2014; Egan et al., 2007, 2010; Enisman et al., 2021; Greenberg & Spiller, 2016; Hagège et al., 2018; Ito et al., 2019; Izuma et al., 2010, 2015; Johansson et al., 2014; Koster et al., 2015; Lieberman et al., 2001; Luo & Yu, 2017; Miyagi et al., 2017; Nakamura & Kawabata, 2013; Salti et al., 2014; Sharot et al., 2009, 2010, 2012; Shultz et al., 1999; Tandetnik et al., 2021; Taya et al., 2014; Voigt et al., 2017, 2019). This phenomenon entails preferences between options that systematically differ (direction and/or magnitude) before versus after choices are made between pairs of the options. The key measurement variable is the *spreading of alternatives* (SoA), because the value ratings of the choice options are typically spread further apart after choices relative to before (see also the literature on *coherence shifts* and *information distortion*: (Carlson & Russo, 2001; Holyoak & Simon, 1999; Russo et al., 1996, 2008; D. Simon et al., 2001, 2004, 2008)). SoA is a robust phenomenon and is particularly high when choices are difficult (e.g., the difference between option values and/or the certainty about option values is low (Lee & Coricelli, 2020; Lee & Daunizeau, 2020)) or when attribute disparity is high (Lee & Hare, 2023b; Lee & Holyoak, 2021). SoA has also been shown to negatively correlate with RT and positively correlate with confidence (Lee & Coricelli, 2020; Lee & Daunizeau, 2020, 2021; Lee & Hare, 2023b; Lee & Holyoak, 2021, 2023).

To calculate SoA, ratings of individual options are obtained before and after choices are made between pairs of options. SoA for each choice is calculated as the difference between the ratings of the chosen and rejected options after the choice minus the rating difference of those options before the choice (Brehm, 1956). Subjective ratings are understood to be noisy estimates of value, and mere statistical noise can give rise to SoA as well as a positive correlation between choice difficulty and SoA (Alós-Ferrer & Shi, 2015; K. M. Chen & Risen, 2010; Izuma & Murayama, 2013). However, there is likely to be a cognitive source of SoA in addition to any SoA caused by noise, according to an analysis of empirical data from multiple previous studies (Lee & Pezzulo, 2023). In that analysis, the authors showed that examining negative SoA from post-to pre-choice ratings (rather than the standard SoA from pre-to post-choice ratings) confirmed that the post-choice ratings contained more choice-relevant information than did the pre-choice ratings. This would not be the case if SoA were purely a statistical artifact, because then the directionality of time would have no meaning and thus no effect on the relationship of SoA (or negative SoA) with choice.

### Sequential sampling model accounts of choice behavior

Various computational models of simple decisions (e.g., two-alternative forced-choice) exist, the most common of which come from the sequential sampling / accumulation-to-bound class (Brown & Heathcote, 2008; Busemeyer & Townsend, 1993; Ratcliff et al., 2016; Usher & Mcclelland, 2001). Under this type of model, the agent processes information about the choice options incrementally until the evidence in favor of one of the options passes a threshold and that option is declared to be the winner (and is therefore chosen). Although multiple variations of this core model class have been proposed, they generally rely on a relative-value accumulator that tallies information about the options based on random samples from underlying probability distributions (one for each option) until it reaches a preset evidence threshold / response boundary. This class of model has provided an elegant account of choice and RT patterns in a variety of domains, in particular with respect to the impact of choice difficulty (value proximity; see (Lee & Usher, 2023) for a version that also includes value uncertainty as a component of choice difficulty). With some minor modifications, this type of model has also simultaneously accounted for the impact of overall set value on choice behavior (Krajbich et al., 2010; Lee & Usher, 2023; Shevlin et al., 2022; Shevlin & Krajbich, 2021; S. M. Smith & Krajbich, 2019; Ting & Gluth, 2025). A multi-attribute version of the model accounts for the impact of attribute disparity (Lee & Hare, 2023a). Even choice confidence can be accounted for by some variations of sequential sampling model, either as a secondary readout of the evidence accumulation process (Calder-Travis et al., 2021; De Martino et al., 2013; Kiani et al., 2014; Moran et al., 2015; Moreno-Bote, 2010; Pleskac & Busemeyer, 2010; Ratcliff & Starns, 2009; van den Berg et al., 2016; Vickers & Packer, 1982; Zylberberg et al., 2012) or as the primary readout itself (Lee, Daunizeau, et al., 2023).

### Sequential sampling models and the spreading of alternatives

The evidence accumulation-to-bound framework has proven to be a powerful tool, yet there remains an important gap in its explanatory power. Despite the success that it has had in accounting for most relevant variables in the study of simple decisions (i.e., accuracy/consistency, RT, and sometimes confidence), it remains to be tested whether it can account for the SoA phenomenon. In its basic form, this is unlikely, by virtue of the very assumptions that it is built on. In brief, sequential sampling models tend to rely on stationary inputs to the evidence accumulation process. For subjective value-based decisions, these inputs are typically regarded to be the modes of latent probability distributions representing the value of each option in the mind of the decision maker. The fact that these inputs are assumed to be stationary precludes the possibility that such traditional models would be used to account for changes in value estimates.

Recent interest has arisen in finding a way to address SoA with sequential sampling models (Lee & Daunizeau, 2021; Zylberberg et al., 2024). Lee and Daunizeau suggest that SoA occurs as a result of a tradeoff between mental effort and choice confidence (Lee & Daunizeau, 2020, 2021). This follows the *resource-rational* approach, where cognition relies on an optimal usage of limited resources (Lieder & Griffiths, 2020). When faced with a difficult decision (e.g., if the options at first seem equally valuable and thus initial confidence about knowing the best option is low), a decision maker might prefer to invest mental effort towards processing additional information with the intention to distinguish the option values more confidently, rather than making an immediate choice with low confidence (Pezzulo et al., 2013; Stoianov et al., 2018). The cognitive processes that eventually result in SoA would therefore be instrumental to the decision as it unfolds, since they would effectively increase the discriminability of the options and make the choice easier. To address SoA, this proposal removes a key assumption of classical sequential sampling models — specifically, that preference responses develop based on value-related signals that are *retrieved* (e.g., from memory) and compared rather than being *constructed* during deliberation (see the literature on *constructed preferences*: (Ariely et al., 2006; DeKay et al., 2011; Lichtenstein & Slovic, 2006; Payne et al., 1999; D. Simon et al., 2008; Tversky et al., 1990; Warren et al., 2011)).

Certain Bayesian models (Li & Ma, 2021; Tajima et al., 2016) contending that evidence accumulation summarizes an updating of option value estimates and certainty could offer a different explanation of SoA (as prior estimates evolve to posterior estimates). Given the known equivalence between Bayesian models and sequential sampling models (Bitzer et al., 2014), it might be interesting to consider the Bayesian approach when seeking a sequential sampling approach. The relevant models assume that both options are initialized with null (or at least equal, if there is prior knowledge about the summary statistics of the population of potential options) prior value estimates, but that the value estimates can come to differ as they are elucidated over time. From this perspective, preferences are not constructed during deliberation and do not change over time; they are simply more clearly recognized as their signals stabilize. This could result in an observed SoA effect between the start and end of deliberation. However, empirical SoA is calculated using subjective value ratings obtained both prior to and after the actual choice. It is unclear how these Bayesian models might account for ratings that were systematically different when elicited after versus before the choice.

Furthermore, such Bayesian models would hold that value estimates evolving during choice deliberation could never shift closer together from start to end of deliberation, since they are initialized identically. Empirically, value estimates often do shift closer together, as evidenced by negative SoA on individual trials (Lee & Holyoak, 2021, 2023). Importantly, this would also mean that decision makers could never change their minds between start and end of deliberation, because all potential preferences would be null at decision onset.^1^ Previous studies have shown such changes of mind (i.e., choices in favor of options that were initially rated lower, but later rated higher than the alternatives) to be common (Lee & Daunizeau, 2020, 2021; Lee & Holyoak, 2023), suggesting that deliberation might sometimes cause an apparent preference reversal if the agent considers new information that it initially neglected (or weighs information differently than it initially did; (Lichtenstein & Slovic, 2006; Tversky et al., 1990)). In standard accumulation-to-bound models, the process starts off with total ambivalence about preference, precluding the very possibility of a change of mind / preference reversal during deliberation.

One potential way to resolve this issue and enable this class of model to provide a measure of SoA would be to allow each option to enter deliberation with its own specific prior value estimate and precision. Mathematically, this would entail initializing the evidence variable at a point of non-neutrality. Previous work has presented sequential sampling models with such a starting point bias, representing prior beliefs about which option is *a priori* more likely to be the better one, based on instruction (Mulder et al., 2012), choice history (Urai et al., 2019), category preference (Lopez-Persem et al., 2016), personality trait (F. Chen & Krajbich, 2018), or choice history during embodied decisions (Kane et al., 2023; Lepora & Pezzulo, 2015; Molano-Mazón et al., 2024; Priorelli et al., 2024). A starting point bias could also be used to capture option-specific priors (Lee, Daunizeau, et al., 2023). These priors would form immediately upon presentation of the choice options, during the time it takes for perceptual processing to inform the decision apparatus about the identities of the options, but before explicit and intentional value deliberation begins. This would align with research showing that value signals in the brain arise automatically (Lebreton et al., 2009), even when not relevant (e.g., during perception). The formation of priors in this way would not only explain the starting point bias, but also a portion of the non-decision time (NDT), which is typically included in sequential sampling models to represent the time required for stimulus encoding or visual pre-processing as well as motor response execution (Fontanesi et al., 2019; Mulder & Maanen, 2013; Nunez et al., 2017). Once the priors were established, the evidence accumulation process at the core of the model would begin, terminating when the desired evidence level was reached. The difference in the posterior value estimates of the options (i.e., those at the end of deliberation based on total accumulated evidence) could then be compared to the difference in the priors to calculate a measure of SoA. In this way, this type of model could also account for changes of mind, because the chosen option would not always be the option with the higher prior value estimate.^2^

The mechanisms presented above would result in positive SoA on average (Lee & Daunizeau, 2021), and they would also account for the robust relationship between choice difficulty (inverse absolute difference in value estimates, adjusted by value certainty) and SoA observed in the empirical data. But if deliberation were formalized as a process of recognition (or purification) of pre-existing value representations (Ratcliff, 1978), the signals that inform the initial and final preferences would have identical means – only the precision would differ. So, any SoA that might arise would be purely due to noise, which does not seem to be the case for empirical observations of SoA (Lee & Pezzulo, 2023).

A more satisfactory way to account for the SoA data would be to focus on the crucial distinction between easy decisions (during which choices largely reflect prior value estimates and hence there is little or no SoA) and difficult decisions (during which novel sources of evidence are considered and hence there is often greater SoA). Choice difficulty might then be defined not based on some comprehensive measure of value but rather on an incomplete measure of value based on whatever (presumably non-exhaustive) information was considered during the initial isolated rating task. Under this account, the intra-decision dynamics of evidence accumulation (e.g., the evolution of the momentary drift rate) would differ as a function of choice difficulty. During easy decisions, when the choice is between options with very different values, evidence based only on the priors might already surpass the response threshold, in which case the choice could be made immediately without the need for deliberation, or with little additional evidence (see (Alós-Ferrer, 2018; Caplin & Martin, 2016; Diederich & Trueblood, 2018; Pezzulo et al., 2013) for alternative models that allow for this sort of automatic choice)^3^. This would imply no meaningful differences between the initial and final preferences and hence no SoA for such easy decisions. Difficult decisions, in contrast, are alleged to elicit higher levels of information processing (Lee & Daunizeau, 2020, 2021; Pezzulo et al., 2013), which would potentially alter the means of the option value representations if newly-considered information were not fully congruent with previously-considered information.

It could be the case that early information systematically differs from late information. For example, it has been shown that decision-relevant information processing related to more salient or important attributes begins earlier than that related to less salient or important attributes (Lim et al., 2018; Maier et al., 2020; N. Sullivan et al., 2015). Furthermore, it has been proposed that the information considered in the formation of prior values and during deliberation is distinct and develops by means of passive retrieval from memory or active sampling via mental simulation, respectively (Pezzulo et al., 2013). Because the information considered at the start and at the end of a decision will not necessarily be the same (and thus the standard assumption of sampling from stationary inputs is removed), the value estimate for each option could change over the course of deliberation. This suggests that the evidence accumulation rate in sequential sampling models should perhaps not be taken as static, but rather dynamic across deliberation time. This motivates the use of an extension to standard models in the form of an intra-trial time-varying drift rate. This approach has previously been suggested using drift rates that alternate or change in discrete stages within each trial (Diederich, 1997, 2003; Diederich & Oswald, 2014; Krajbich et al., 2010; Maier et al., 2020) or continuously over time (P. L. Smith, 2000). One recent proposal has even suggested that noise in the evidence accumulation process gets absorbed as true evidence, giving rise to a drift rate that continuously changes throughout a decision (Zylberberg et al., 2024).

In this study, we examine the classic drift-diffusion model under our assumptions outlined above regarding how SoA might arise during the sampling and accumulation process. We first examine some simple variants of the model and validate their fundamental capability to produce SoA in general. We then test to determine which modifications might be necessary to produce the SoA effects previously reported in the literature.

## Methods

Using simulation and regression analysis, we test whether standard or modified drift-diffusion models (DDMs) can account for previous findings related to the spreading of alternatives (SoA) effect. We based our simulations on experimental data pooled together from two previous studies (n = 580; (Lee & Hare, 2023b; Lee & Holyoak, 2021)). In those studies, participants performed modified versions of the free-choice paradigm, where they first provided isolated ratings for each member of a set of options (snack foods), then chose between pairs created from the same set of options (each option appeared in only one pair), then again provided isolated ratings for each option. The tasks were modified from the standard free-choice paradigm in that participants here provided ratings for both overall value (as always) and individual attributes (pleasure and nutrition) for each option. We excluded participants who seemed to not perform the tasks correctly, which we defined as having a slope coefficient of less than one in a logistic regression of choice on value ratings (n = 17). There was also a technical issue in one of the original experiments where the data for one participant did not save properly, leaving us with n = 562 on which to conduct our analyses. For all participants, we excluded trials where RT was either lesser than 0.5 seconds or greater than 10 seconds, as is standard in this type of research.

The dataset we worked with thus included all the variables we needed for our simulations. Specifically, we used as input to our simulations the ratings that each participant provided for the overall value (*V*), the first attribute (pleasure or *P*), and the second attribute (nutrition or *N*) of each option, as well as the pairings of left and right options that each participant encountered on each choice trial. We calculated the difference in overall value (*dV*) for each choice trial as the difference between the ratings of overall value (option 1 minus option 2). We calculated the differences in the first and second attributes (*dP* and *dN*, respectively) in the same manner. We used the estimated coefficients from a regression of overall value on the attributes (separately for each participant) as the participant-specific attribute weights needed to calculate attribute disparity (orthogonal to dV) for each trial, as defined by Lee and Holyoak (Lee & Holyoak, 2021):

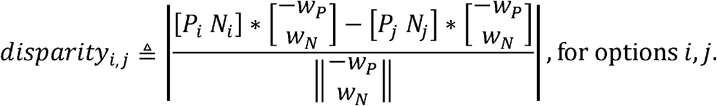

where P and N refer to the ratings for the first and second attributes, respectively, and *w* refers to the weight of the contribution of each attribute to overall value.

Next, we simulated the evidence accumulation process for each trial for each participant (under the basic DDM as well as each of the modified versions we examined; details about how the evidence accumulation process proceeded under each model formulation are provided in Appendix A.) For the model parameters, we used the best-fitting parameters for each participant obtained from fitting the experimental choice and response time data using the experimental rating data for value, pleasure, and nutrition (details of our model-fitting procedure are provided in Appendix B.) At the onset of each trial, time and net evidence were initialized to zero. With each time step (*t*) beginning after the non-decision time, an increment of net evidence was added to the cumulative net evidence. The process stopped when the magnitude of the cumulative net evidence reached the response threshold. The choice response (*Ch*) on each trial was recorded as option 1 if the final cumulative net evidence (*E*) was positive or option 2 if E was negative (coded as 1 for option 1, 0 for option 2). RT was recorded as the final time step (*T*) on each trial plus non-decision time (NDT; a free parameter of the models), divided by 1,000 to make the quantities representative of seconds. Spreading of alternatives (SoA) on each trial was calculated as follows:

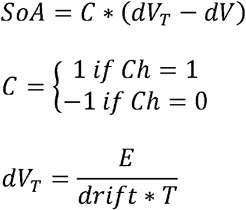

In this way, dV_T_ can be thought of as the “inferred” value difference based on total accumulated evidence and deliberation time^4^, whereas dV can be thought of as the “anticipated” value difference based on previous evaluations. The formula for SoA includes C because dV is calculated with respect to option 1 minus option 2, but SoA is defined with respect to the chosen option minus the rejected option.

After creating the simulated data, we tested for relationships between the relevant decision variables: |dV| (as a measure of difficulty, with smaller values indicating greater difficulty), disparity (D), consistency, RT, and SoA. (As with the experimental data, we excluded trials where RT was either lesser than 0.5 seconds or greater than 10 seconds.) Specifically, we ran a series of regression analyses in Matlab using the *fitlme* function for the linear regression models and the *fitglme* with a binomial distribution and logit link for the logistic regression models. We included random effects (both slopes and intercepts) for each participant in all regression models. For comparison with the simulated results, we ran the same series of regressions using the experimental data from previous studies (Lee & Hare, 2023b; Lee & Holyoak, 2021). Note that one of the Lee & Hare experiments did not include post-choice ratings, so the data from those participants (n = 107) were not included in any of the SoA analyses.

## Results

### Confirmatory results

As an initial check that our simulated data sufficiently resembled the experimental data, we examined the degree to which choices were consistent with reported value ratings (consistency, averaged across participants). For difficult choices (defined as having |dV| below the median of 0.21), the average consistency was 68% in the experimental data. For easy choices (|dV| above the median), the average consistency was 92% in the experimental data. We also examined response times. For difficult choices, the median RT was 1.8s in the experimental data. For easy choices, the median RT was 1.6s in the experimental data. In the data simulated under the DDM variants, difficult choices had an average consistency of 55-60% and an average RT of 1.9-2.0s; easy choices had an average consistency of 81-91% and an average RT of 1.5-1.7s. For a more refined perspective on how well the different model variants reproduced the experimental choice data, we created quantile plots of consistency across bins of |dV| median split by fast and slow RT, and across bins of RT median split by easy and difficult |dV| (note: these median splits are on the entire data table, not separately within bins). Figure 1 illustrates how well the patterns in the simulated data matched those in the empirical data. Note that all model variants do a reasonable job of replicating the patterns.

**Figure 1:**
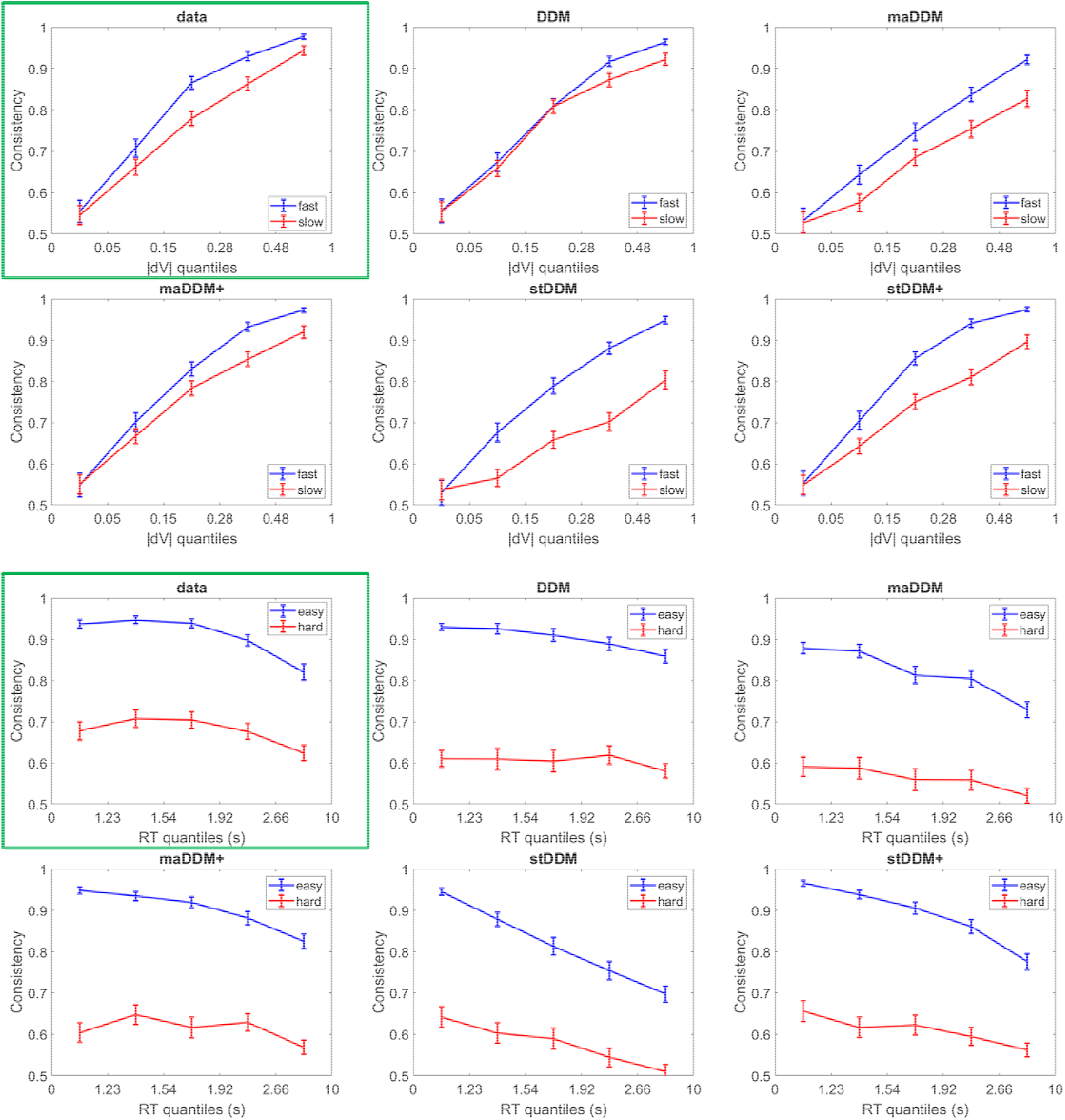
Quantile plots for choice consistency as a function of difficulty and RT. All data was binned based on the empirical dataset. The median splits were based on the full empirical dataset.

Previous studies reported that choice consistency had a positive relationship with both |dV| and D and that RT had a negative relationship with both |dV| and D (Bhatia & Mullett, 2018; Lee & Hare, 2023b; Lee & Holyoak, 2021). We thus regressed consistency (logistic) and log(RT) separately on both |dV| and D, using the experimental data as well as the data simulated under the DDM variants. We replicated the previous findings in the experimental data. The simulated data qualitatively reproduced all patterns with respect to |dV|. With respect to D, results differed across the model variants. The basic DDM was unable to produce any relationship between D and either dependent variable, due to its ignorance of attributes. Variants that included attribute ratings, however, were able to qualitatively replicate some of results. The maDDM and stDDM reproduced the positive relationship between D and consistency, and all multi-attribute variants reproduced the negative relationship between D and RT. (See Figure 2.)

**Figure 2:**
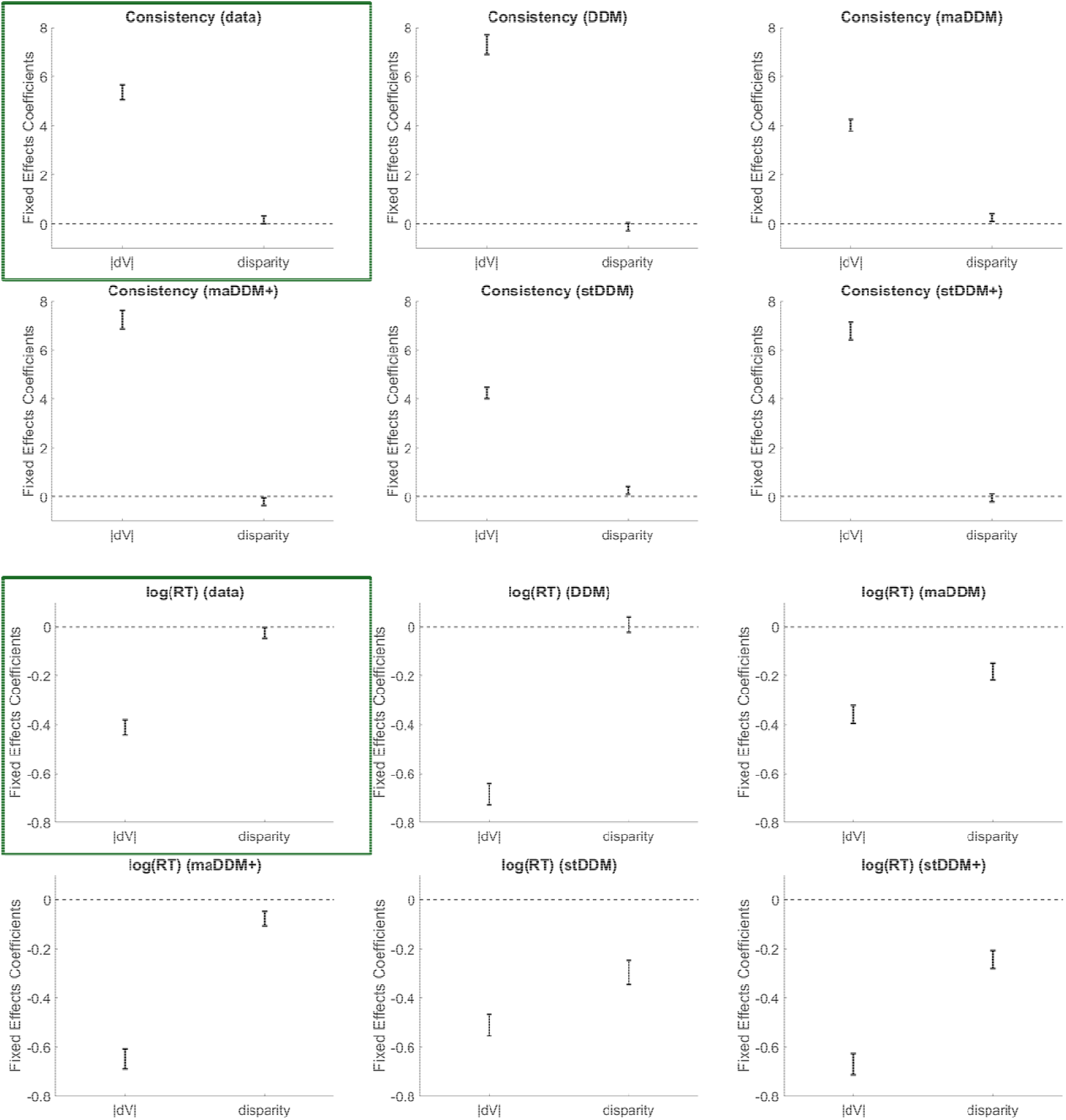
Regression coefficients for choice difficulty (|dV|) and attribute disparity on consistency and RT. Error bars represent 95% confidence intervals.

We next checked whether the models were able to output a positive average SoA acros trials, and especially for trials on which the decision was difficult. For easy and difficult choices, the mean SoA was -0.02 and 0.08, respectively, in the experimental data. In the data simulated under the DDM variants, SoA ranged from -0.13 to 0.15 for easy choices and from 0.18 to 0.40 for difficult choices. Each of the model variants output positive SoA across all trials and especially for difficult trials, although the magnitudes SoA varied substantially across models. This could suggest that certain model variants better capture the effect, or that the way in which we calculated SoA needs to be more precisely tuned, possibly differently for each model. For a more refined perspective on how well the different model variants reproduced the experimental SoA patterns, we created quantile plots of SoA across bins of |dV| median split by fast and slow RT, and across bins of RT median split by easy and difficult |dV|. Figure 3 illustrates how well the SoA patterns in the simulated data qualitatively matched those in the empirical data. Note that all model variants reproduced the general qualitative patterns, although some do a better quantitative job than others as noted above).

**Figure 3:**
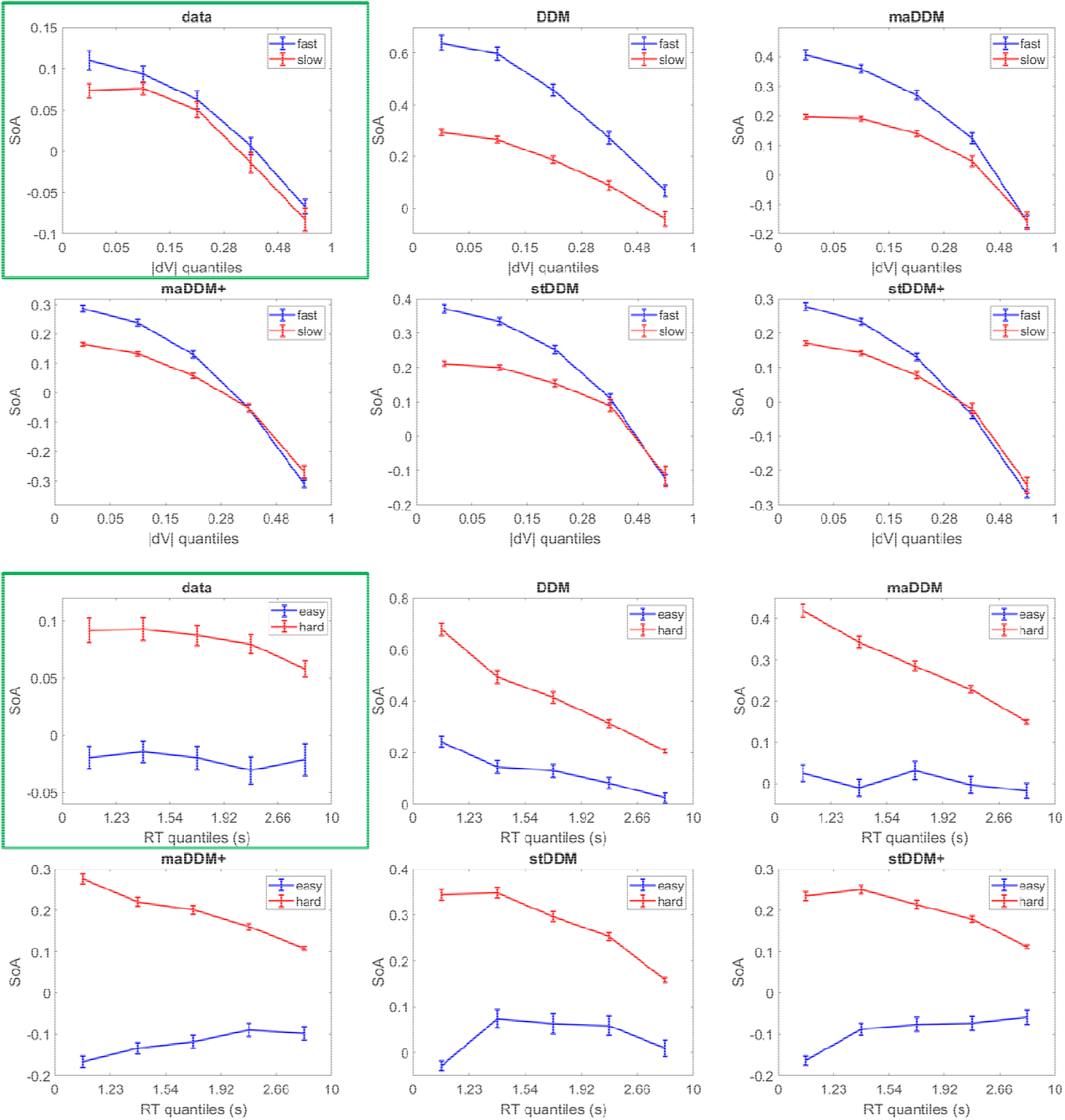
Quantile plots for spreading of alternatives as a function of difficulty and RT. All data was binned based on the empirical dataset. The median splits were based on the full empirical dataset.

Previous studies reported a negative relationship between |dV| and SoA and a positive relationship between D and SoA (Lee & Hare, 2023b; Lee & Holyoak, 2021). We thus regressed SoA on |dV| and D, using both the experimental and the simulated data. We replicated the previous findings in the experimental data. The simulated data qualitatively reproduced the pattern with respect to |dV|. This is not surprising, given previous reports about how SoA can arise from mere statistical noise (Alós-Ferrer & Shi, 2015; K. M. Chen & Risen, 2010; Izuma & Murayama, 2013; Lee & Pezzulo, 2023). With respect to D, results differed across the model variants. The basic DDM did not produce any relationship between D and SoA, but variants that included attribute ratings qualitatively replicated the patterns of effects (Figure 4). Previous studies also reported that SoA had a clear negative relationship with RT that is qualitatively similar to the relationship between |dV| and RT (Lee & Hare, 2023b; Lee & Holyoak, 2021, 2023; Lee & Pezzulo, 2023). We thus tested for this as well, regressing log(RT) on |dV| and SoA. We replicated the previous findings in the experimental data. The data simulated under all DDM variants qualitatively reproduced the pattern of effects (Figure 4). This is not surprising, as SoA is equivalent to an increase in drift rate under the DDM (or to the choice becoming easier, in more general terms), and increased drift rates enable faster responses.

**Figure 4:**
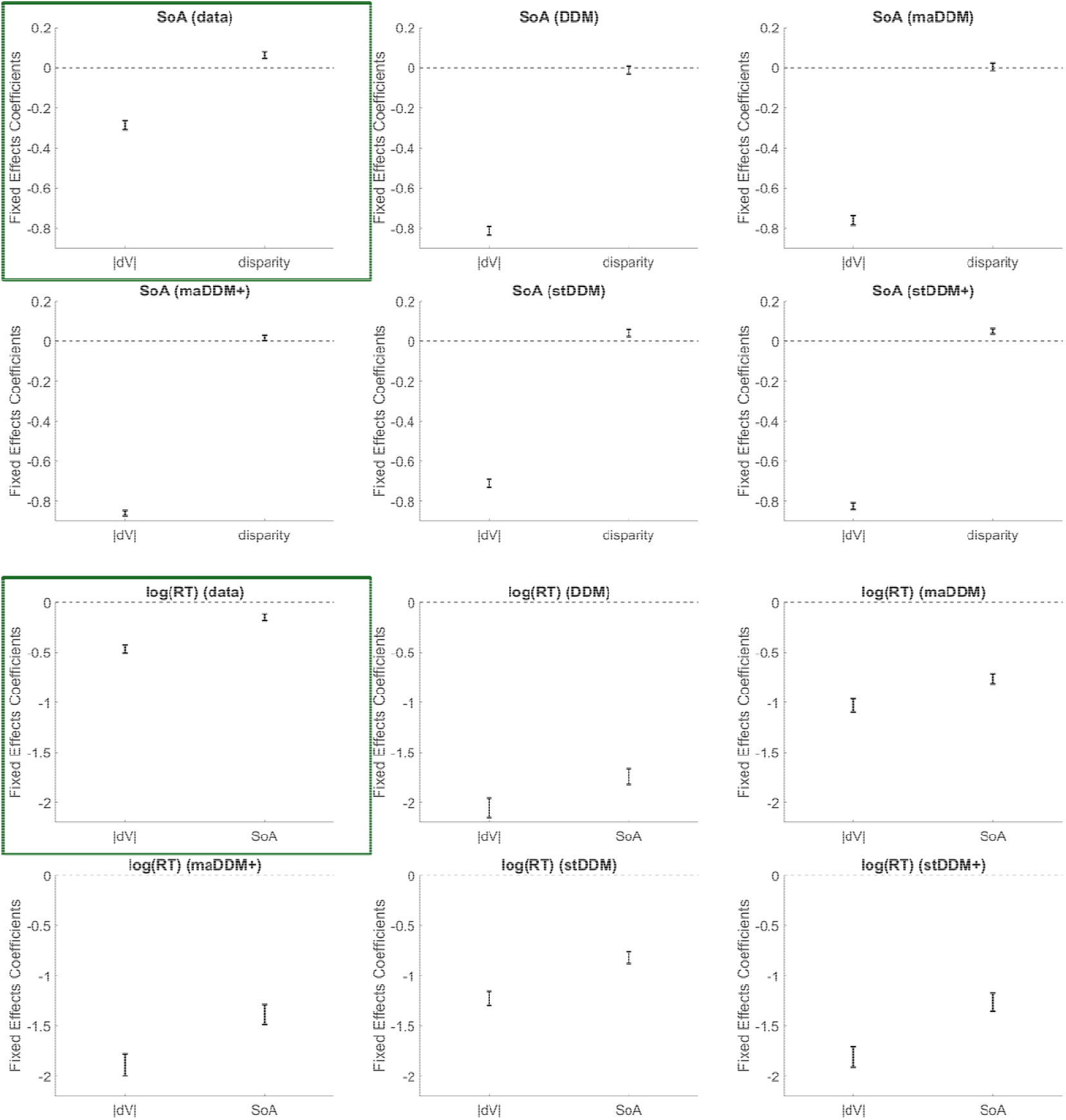
Regression coefficients related to spreading of alternatives (SoA). Error bars represent 95% confidence intervals.

## Discussion

In this work, we investigated whether evidence accumulation-to-bound models of value-based decision-making can account for choice-induced preference change (as measured by the spreading of alternatives or SoA) in addition to response probability and response time (RT), which they have been well-documented to simultaneously account for. We focused on the drift-diffusion model (DDM) because of its simplicity and analytical tractability, and because of its widespread success in accounting for standard decision variables. In addition to the classic DDM, we also considered several variants that included simple extensions. Most of these model variants have been presented in previous work, although none of those studies examined choice-induced preference change. Our results provide an account for how SoA could arise during choice deliberation by including information about individual attributes in the evidence accumulation process and by assuming that value estimates at the end of the decision process are informative about future individual value ratings.

This study establishes proof of concept that the DDM (and presumably, a wide variety of sequential sampling models) can generate SoA in value-based decisions based on multiple attributes. Our results show that the key factor is to include different evidence streams for each attribute. We demonstrate via simulation that this is sufficient to give rise to SoA, complete with the correct (i.e., in line with the empirical findings) relationships between SoA and both choice difficulty and attribute disparity, as well as between RT and SoA. Including different evidence onset latencies for different attributes may help but does not seem to be necessary. Thus, the well-established SoA phenomenon can be fully explained by a principled process-based model, removing the need to rely on more descriptive accounts such as the post-choice resolution of cognitive dissonance (Brehm, 1956; Festinger, 1957).

The mechanistic process we considered provides an account for how SoA between choice options might arise *during* a decision. However, such intra-decisional changes in evaluations are typically not observable (but see (Lee & Holyoak, 2023)). For this reason, experimental evidence for SoA generally comes in the form of isolated evaluations of the choice options, which are (on average) more distant after the choice task relative to before. Under the model described above, one would need to assume that pre-choice evaluations are made based on either a different set of attributes than the set that determines post-choice evaluations or a set of attributes that is differently biased towards a particular subset. For example, in a decision about snack foods, a chocolate bar and an orange both might have been initially (pre-choice) rated very highly (in isolation), based predominantly on their respective tastiness. During a decision in which these options were set against each other, however, the tastiness attribute might not enable a clear choice and so the healthiness attribute might be considered more before the orange is chosen as the best option. Then, when the items are later re-evaluated in isolation (for post-choice ratings), the orange might be rated higher than the chocolate bar, because now the healthiness attribute (which favored the orange) is more influential in the overall ratings than it was before the choice. We focus here on identifiable attributes, but in a more general sense, the idea is that earlier assessments (either in isolation or in tandem) are less information-rich than later assessments, provided that the agent has proper motivation to seek enrichening information (Bénon et al., 2024; Lee & Daunizeau, 2020, 2021). Most likely, the posterior beliefs about option values from one task will inform the prior beliefs for a subsequent related task, following standard Bayesian computations.

When an agent initially considers each option in isolation (e.g., during a pre-choice rating task), different attributes might be more or less salient for different options. In that case, for some options, the value estimates at the start of the decision might relate more to one attribute, whereas for other options, the initial value estimates might relate more to a different attribute. Additionally, when choosing with a specific goal in mind, there could be a default primary attribute that the agent would consider before the others regardless of what the options were (Lim et al., 2018; Maier et al., 2020; N. Sullivan et al., 2015). Such an attribute should always be salient. These ideas combine to suggest that both option-specific factors and task-related factors could come into play to give rise to SoA. Predicting the change in attribute evaluations that might occur would be more complex, as some SoA could cause a particular attribute to have more impact on the choice while other SoA could cause the opposite. A future study dedicated to examining the rating process might help to better explain the intricacies of these various factors.

The ability of the DDM to generate SoA (as we have presented it) relies on the assumption that post-choice ratings will resemble the latent value estimates that are constructed during choice deliberation up until the time a choice is made. This idea would generally align with the idea that value estimates are updated in a Bayesian manner during deliberation (Tajima et al., 2016), where the posterior estimates would endure without decaying back to the priors. It has been shown that the SoA effect is maximal *during* choice deliberation, although it then partially diminishes during a subsequent rating task (Lee & Holyoak, 2023). This suggests that post-choice ratings might be based on some combination or interpolation of the information on which pre-choice ratings were based and the information on which choices were based. It seems that during isolated evaluations after choices have recently been made, the agent will tend to focus more on whichever attributes contributed to the basis of comparison during the relevant choices. It is as if thinking about (or not) attributes during choice deliberation made those attributes more (or less) likely to be activated when considering the options again in the future (even in isolation). This would confer choices some degree of history dependence that is not generally accounted for by current models. Future work should investigate the transient nature of SoA, as well as the neural underpinnings of bases of comparison during pre-choice rating, choice, and post-choice rating tasks.

Most studies related to SoA, including this one, consider only changes in ratings of overall value, but are agnostic about ratings of individual attributes. However, it has also been shown that SoA occurs even at the level of individual attributes (Lee & Holyoak, 2021). In other words, attribute evaluations tend to favor the chosen options more after the choices are made, relative to before (Lee & Holyoak, 2021). Perhaps SoA at the attribute level is caused by the same mechanisms that we have proposed for SoA at the level of overall value. This could be formalized as weights of sub-attributes (onto attributes and subsequently onto overall value) that differ for choice versus rating tasks. Note that certain contextual aspects of value-based decision making could be related to higher-level attributes being broken down into sub-attributes in the mind of the agent (e.g., when choosing a car, size could merely be a proxy attribute for the actual attributes, which might include comfort, utility, capacity, convenience, and ease of use). So, it could be said that deliberation involves not only the consideration of different attributes, but also different sub-attributes (in the DDM, these could also be represented as different evidence streams with unique drift rates). Future work could explore sub-attributes and in what ways their relationship to higher-level attributes mirrors the relationship of attributes to overall value. This might build upon previous work showing that describing an attribute in different ways to highlight different objectives related to that attribute can alter preference judgments (Mertens et al., 2020; Ungemach et al., 2018).

## Code Availability

The analysis code used in preparation of this manuscript is publicly available on the Open Science Framework at https://osf.io/2bs73/.

## Appendix A

Computational models

### 1. Drift-diffusion model (DDM)

The first model we considered was a classic standard DDM, governed by the following equations:

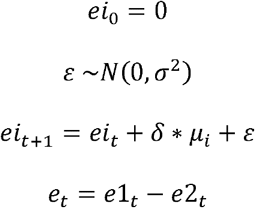

Where *ei*_*t*_ is the momentary evidence for option *i* ∈ {1,2}, *t* is an index for deliberation time (arbitrary units), ε is the white processing noise with strength *σ*^2^ common to every decision, *μ*_*i*_ is the value estimate of option *i, δ* is an agent-specific scalar representing evidence efficiency, and *e*_*t*_ is the momentary differential evidence in favor of option 1 over option 2.

### 2. Multi-attribute drift diffusion model (maDDM)

The second model we considered was a multi-attribute version of the DDM (maDDM; (Lee & Hare, 2023a)). Expanding the basic DDM so that the process is based on multiple attributes is straightforward, replacing the overall value estimates *μ*_*i*_ with estimates of individual attributes 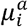 and the drift scalar *δ* with a vector of attribute-specific drift scalars *δ*^*a*^ for each attribute *a*. The equations governing this model are:

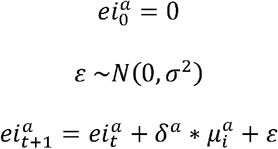

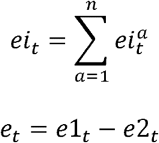

where *n* is the dimensionality of the attribute space for the decision at hand. Note that the scaling vector *δ*^*a*^ is mathematically equivalent (and notationally simpler) to a vector of attribute weights multiplied by a general scaling factor specific to the agent and common across attributes.

### 3. Multi-attribute plus value drift diffusion model (maDDM+)

The third model we considered was a hybrid of the maDDM and the standard DDM (maDDM+; (Lee & Hare, 2023a)). This model was previously shown to dominate both of the component models in a formal model comparison (Lee & Hare, 2023a). The equations governing this model are the same as those for the maDDM, with overall value entering the equations as if it were a third attribute.

### 4. Start-time drift diffusion model (stDDM)

The fourth model we considered was one in which evidence for different individual attributes entered the accumulation process at different times, making it a generalization of the maDDM. This model was previously shown to account well for attribute-based decisions and to dominate the maDDM in a formal model comparison (Maier et al., 2020; N. J. Sullivan & Huettel, 2021). The equations governing this model are the same as those for the maDDM, except that *δ*^*a*^ = 0 ∀ *t* < *S*^*a*^, where *s* is the start time for each attribute to enter consideration.

### 5. Start-time plus value drift diffusion model (stDDM+)

The fifth model we considered was a hybrid of the stDDM and the standard DDM (stDDM+). The equations governing this model are the same as those for the stDDM, with overall value entering the equations as if it were a third attribute. Evidence for overall value always starts accumulating before evidence for either of the individual attributes.

## Appendix B

Model-fitting procedure

We fit each of the candidate models to the experimental data in order to recover the best parameters for each model to explain the data. For this model fitting exercise, we relied on the Variational Bayesian Analysis toolbox (VBA, available freely at https://mbbteam.github.io/VBA-toolbox/; (Daunizeau et al., 2014)) with MATLAB R2024a. Within participant and across trials, we entered the ratings of overall value, pleasure, and nutrition for each option as input and choice and RT as output. We also entered the model-specific mappings from input to output as outlined in the analytical formulas in the main text. We fixed the diffusion noise (σ^2^) to 0.5, so the parameters to be fitted were the drift scalar (*d*), response threshold (θ), non-decision time (NDT), and start time (*s*^*a*^) terms described above in the model formulations. VBA requires prior estimates for the free parameters, which we set to 1 for *d* and θ (constrained to be positive) and 0.4 for NDT and *s*^*a*^ (constrained to be between 0 and 1). VBA then recovers an approximation to the posterior density on unknown variables. We used the VBA_NLStateSpaceModel function to fit the data for each participant individually.

VBA estimates parameters during model fitting using Variational Bayes: an efficient iterative algorithm that provides a free-energy approximation for the model evidence, which trades off model accuracy (goodness of fit, or log likelihood) and complexity (degrees of freedom, or KL divergence between priors and fitted parameter estimates; see (Friston et al., 2007; Penny, 2012)). The VBA algorithm starts with our relatively flat Gaussian priors for each model’s free parameters and eventually provides a posterior density estimate. Previous studies have successfully used this approach to fitting variants of DDM (Feltgen & Daunizeau, 2021; Lee, D’Alessandro, et al., 2023; Lee & Hare, 2023a; Lee & Usher, 2023; Lopez-Persem et al., 2016).

## Footnotes

1 This statement would no longer hold if Bayesian models were augmented with option-specific prior values (Tajima et al., 2016).

2 One further assumption that this approach would require is that the process resulting in initial isolated value estimates for each option (during the pre-choice rating task in a typical experimental paradigm) is similar to that which establishes the initial imprecise value representation at the start of a decision (because SoA is measured experimentally based on pre-choice ratings, as any latent value representations that form at the beginning of a choice are not observable). Although this assumption might seem unreasonable, it may nevertheless be reasonable to assume that the process that yields isolated ratings lies somewhere between the very imprecise prior estimates and the relatively precise posterior estimates (i.e., it is a similar process but with an intermediate level of precision).

3 Alternatively, in tasks where participants are required to report a decision by reaching or clicking with the computer mouse one of two response buttons located far away from the “start” position, the prior could determine the initial direction of (finger or mouse) movement, which may eventually be revised online during deliberation (Barca & Pezzulo, 2012, 2012; Dotan et al., 2018; Lee, D’Alessandro, et al., 2023; Lepora & Pezzulo, 2015).

4 This is not to suggest that the new value estimate is calculated by the agent only after deliberation has concluded. Rather, the state of the evidence accumulator as a function of time can be thought of as the momentary estimate of the value difference at any point in the decision process. The final estimate (dV_T_) is special only because it is what the agent will register as its best estimate, after which no further updates will be expected.

